# CRISPR screen identifies BAP1 as a deubiquitinase regulating SPIN4 stability

**DOI:** 10.1101/2025.08.14.670297

**Authors:** Alondra Sanchez, Chen Zhou, Rima Tulaiha, Francisco Ramirez, Lu Wang, Xiaoyu Zhang

**Affiliations:** Department of Chemistry, Northwestern University, Evanston, Illinois 60208, United States; Department of Biochemistry and Molecular Genetics, Northwestern University, Chicago, Illinois 60611, United States; Simpson Querrey Center for Epigenetics, Northwestern University, Chicago, Illinois 60611, United States; Chemistry of Life Processes Institute, Northwestern University, Evanston, Illinois 60208, United States; Robert H. Lurie Comprehensive Cancer Center, Northwestern University, Chicago, Illinois 60611, United States; Center for Human Immunobiology, Northwestern University, Chicago, Illinois 60611, United States; International Institute for Nanotechnology, Northwestern University, Evanston, Illinois 60208, United States

## Abstract

Protein homeostasis is tightly controlled by the coordinated actions of E3 ubiquitin ligases and deubiquitinases (DUBs). We identify Spindlin-4 (SPIN4), a histone H3K4me3 reader, as a substrate regulated by opposing pathways: degradation mediated by the Cullin-RING E3 ligase DCAF16 and stabilization by the DUB BAP1. Through CRISPR-Cas9 knockout screens and biochemical analyses, we demonstrate that DCAF16 promotes SPIN4 degradation, while BAP1 interacts with and stabilizes SPIN4 through its catalytic activity. Inhibition or loss of BAP1 reduces SPIN4 levels, highlighting its critical role in maintaining SPIN4 homeostasis. Proteomics and interactome analyses further support this regulatory axis. These findings reveal a dynamic balance controlling SPIN4 stability, with potential implications for epigenetic regulation and disease processes.

## Introduction

The ubiquitin-proteasome system (UPS) is a central mechanism for regulating protein turnover and quality control in eukaryotic cells, playing a critical role in maintaining protein homeostasis and orchestrating a wide range of cellular processes^1, 2^. It mediates the selective removal of proteins through the covalent attachment of ubiquitin, a conserved 76-amino acid polypeptide, to lysine residues on substrate proteins, marking them for degradation by the proteasome^3^. The ubiquitination process is driven by a cascade of E1 activating enzymes, E2 conjugating enzymes, and E3 ubiquitin ligases, the latter of which determine substrate specificity^4^. More than 600 E3 ligases are encoded in the human genome, with RING-type ligases comprising the largest and most diverse subclass^5^. Counterbalancing the activity of E3 ligases are deubiquitinases (DUBs), a family of proteases that remove ubiquitin from target proteins or process polyubiquitin chains^6, 7^. DUBs play essential roles in editing or reversing ubiquitination signals, thereby regulating protein stability and signaling pathways^8^. Their dynamic interplay with E3 ligases ensures tight control over the fate of ubiquitinated proteins and contributes to the broader regulation of proteostasis.

We previously identified DCAF16, a Cullin-RING E3 ubiquitin ligase (CRL), while developing electrophilic Proteolysis Targeting Chimeras (PROTACs) for targeted protein degradation^9^. As DCAF16 remains poorly characterized with undefined physiological roles, we further employed a proteomics-based approach to uncover its native substrates and identified Spindlin-4 (SPIN4), a reader of histone H3 trimethylated at lysine 4 (H3K4me3), as an endogenous target of DCAF16^10^. SPIN4 is also poorly characterized but is thought to be involved in epigenetic regulation through its Tudor domain, which binds methylated histones. Only a few studies have explored SPIN4’s potential physiological functions, including its loss-of-function causing X-linked overgrowth syndrome^11^ and its high expression being associated with advanced nodal status^12^. Although our initial study revealed aspects of the SPIN4 regulatory pathway, key questions remain: Is DCAF16 the sole E3 ligase responsible for regulating SPIN4 turnover? And does a specific DUB counterbalance DCAF16’s activity to maintain SPIN4 homeostasis? Addressing these questions is critical for establishing a mechanistic framework to understand how SPIN4 contributes to both physiological and pathological processes. In this study, we employed functional genomics approaches to systematically characterize the E3 ligase and DUB responsible for modulating SPIN4 homeostasis.

## Results and Discussion

### Establishment of a screening platform to identify regulators of protein stability

Clustered regularly interspaced short palindromic repeats (CRISPR)-based knockout screening approaches have proven effective for identifying E3 ligases that regulate the stability of target proteins, either in their native state or in response to small-molecule degraders^13-15^. A commonly used strategy involves tagging the target protein with a fluorescent reporter, such as green fluorescent protein (GFP). When a CRISPR sgRNA library targeting E3 ligase genes is introduced, loss of an essential ligase can lead to impaired degradation and increased target protein abundance, resulting in elevated GFP intensity. This fluorescence change enables the isolation of affected cells via flow cytometry, followed by next-generation sequencing (NGS) to identify enriched sgRNAs targeting key E3 ligase genes.

We first aimed to establish this screening platform. As a model system, we selected a well-characterized ligand-induced degradation pathway in which the target protein is FK506-binding protein 12 (FKBP12) and the degrader, dFKBP12, is a potent heterobifunctional small molecule that efficiently promotes FKBP12 degradation^9^. This system enables rapid depletion of the target protein, allowing clear distinction between wild-type and E3-deficient cells by flow cytometry. Importantly, dFKBP12 leverages lenalidomide as the E3-recruiting moiety, which has been extensively characterized to engage the E3 ligase cereblon (CRBN)^16, 17^. As such, this system provides a tractable and robust framework to validate the feasibility of using CRISPR-based knockout screening to identify regulators of protein homeostasis.

To construct the reporter system, we cloned FKBP12 into a vector in which it is fused to enhanced GFP (EGFP). This construct also includes a separately translated mCherry reporter that is not affected by FKBP12 degradation, thereby serving as a control for gene expression and cell viability (**Figure 1a**). We generated HEK293T cells stably expressing FKBP12-EGFP and treated them with dFKBP12 (**Figure 1b**). Treatment led to a marked reduction in FKBP12-EGFP fluorescence, while mCherry levels remained unchanged (**Figure 1c**). Co-treatment with MLN4924, a neddylation inhibitor^18^, fully blocked the FKBP12-EGFP degradation, consistent with the requirement of CRL activity for CRBN function^19^.

**Figure 1.**
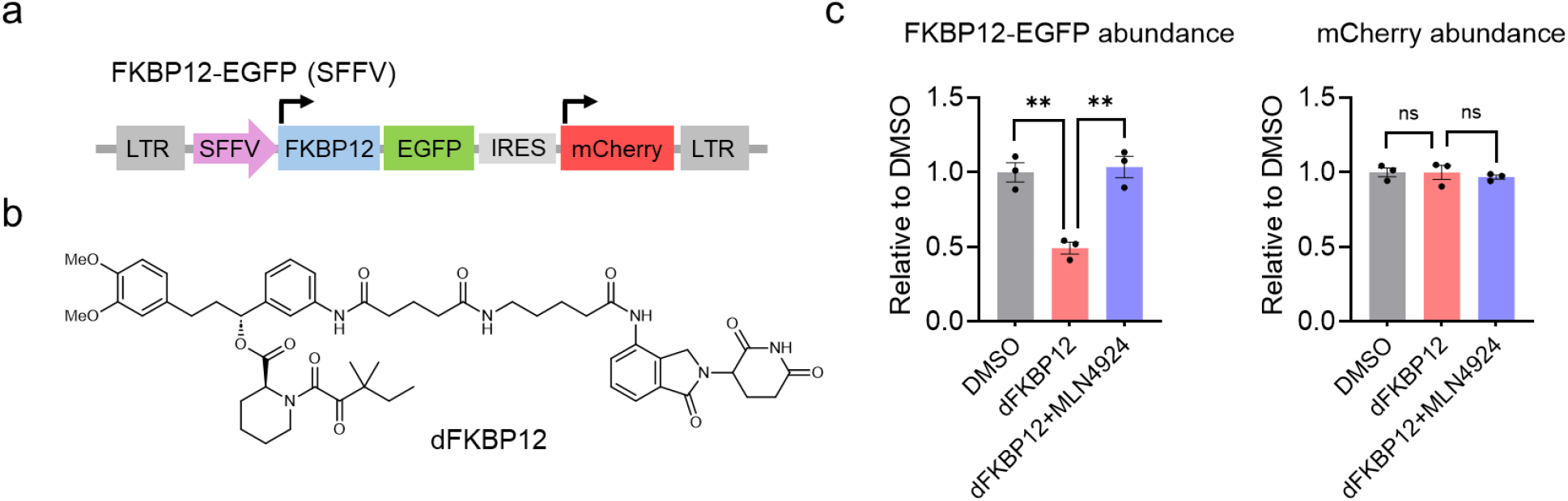
Establishment of an EGFP-based reporter system to study protein degradation. **a**, The construct of FKBP12-EGFP. **b**, Structure of dFKBP12. **c**, Fluorescence quantification of FKBP12-EGFP and mCherry levels in HEK293T cells treated for 8 hours with 2 μM dFKBP12 alone or in combination with 1 μM MLN4924. Data are presented as mean ± SEM (n = 3 biological independent samples). The statistical significance was assessed using unpaired two-tailed Student’s t-tests.

Next, we aimed to perform a CRISPR-Cas9 pooled knockout screen to identify E3 ligases required for dFKBP12-mediated degradation of FKBP12. Cas9 was stably expressed in FKBP12-EGFP-expressing HEK293T cells. A focused sgRNA library was then constructed, containing 4,200 sgRNAs targeting 680 human E3 ligases (six sgRNAs per E3 ligase gene; **Table S1**). To serve as internal quality control, the library also included sgRNAs targeting the EGFP gene. This library was packaged into lentivirus and used to transduce the FKBP12-EGFP/Cas9-expressing cells at a multiplicity of infection (MOI) of 0.3. Following transduction, cells were selected with puromycin and treated with dFKBP12. The top 10% of cells with the highest GFP intensity were isolated via fluorescence-activated cell sorting (FACS) (**Figure 2a** and **Figure S1**). Subsequently, we used NGS to compare the relative abundance of sgRNAs in the sorted cells versus the unsorted input population, identifying sgRNAs that were enriched (**Table S2**).

**Figure 2.**
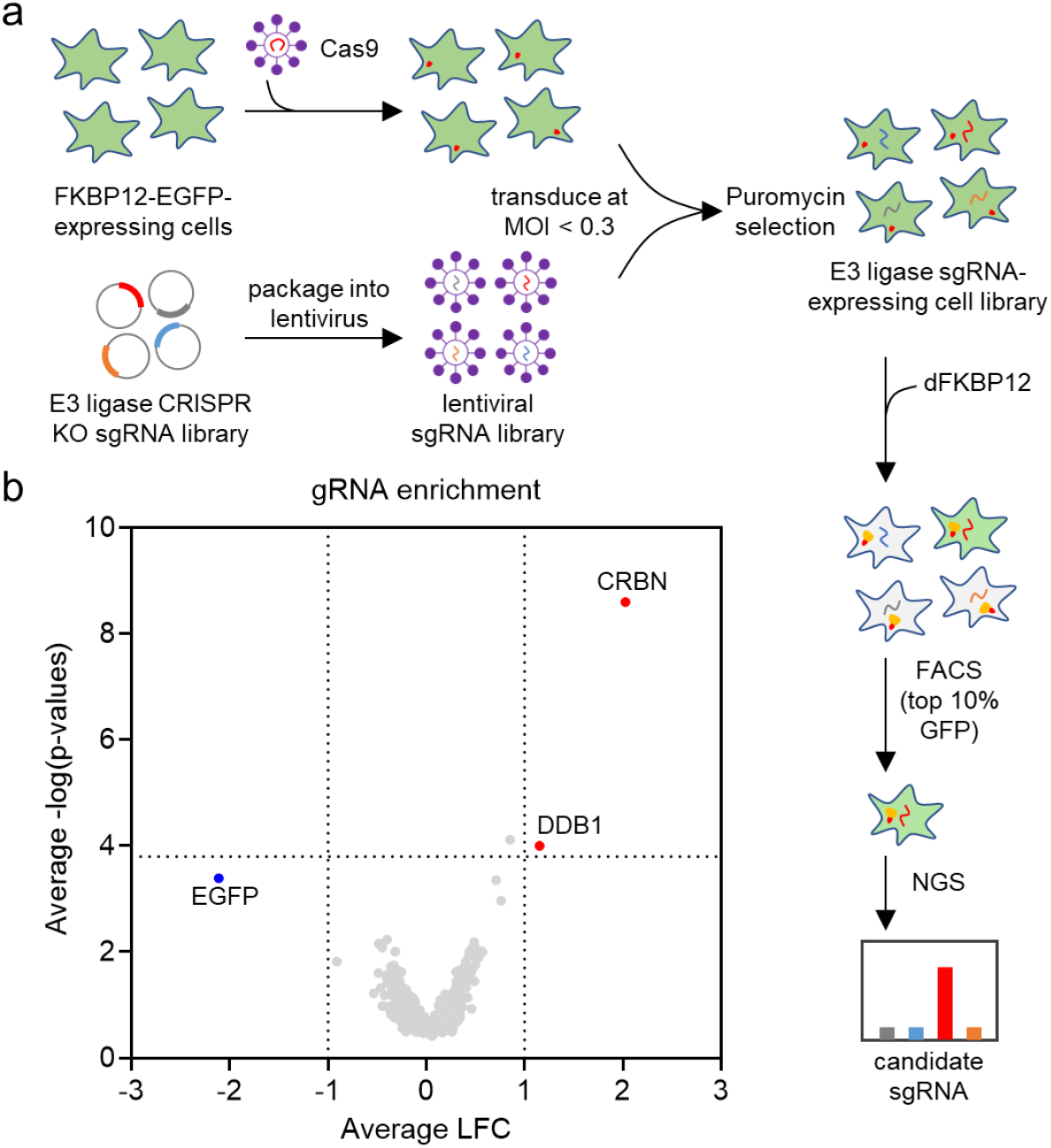
E3 ligase-focused CRISPR-Cas9 knockout screen to identify the E3 ligase required for dFKBP12-mediated FKBP12-EGFP degradation. **a**, Schematic overview of the CRISPR-Cas9 knockout screen workflow. **b**, Volcano plot showing results from the E3 ligase-focused CRISPR-Cas9 knockout screen for FKBP12-EGFP degradation following treatment with 1 μM dFKBP12 for 8 hours (n = 3 biologically independent samples). *P* values were calculated by two-sided t test and adjusted using Benjamini-Hochberg correction for multiple comparisons. LFC, log^2^ fold change.

From this screen, sgRNAs targeting CRBN emerged as the most enriched, indicating that loss of CRBN blocks dFKBP12-induced degradation of FKBP12-EGFP (**Figure 2b**). We also identified DDB1 as a hit (**Figure 2b**), consistent with its known role as an adaptor linking CRBN to the Cullin scaffold, and supporting its essentiality in this degradation pathway. Additionally, sgRNAs targeting EGFP were among the most depleted, as expected, since EGFP knockout would lead to reduced GFP signal and exclusion from the top 10% GFP-intensity population. Collectively, these results demonstrate the effectiveness of our screening platform in identifying regulators of protein stability.

### CRISPR-Cas9 knockout screening reveals DCAF16 as the sole E3 ligase regulating SPIN4 stability

With this screening platform, we aimed to systematically investigate regulators of SPIN4 homeostasis. We first focused on E3 ligases involved in its degradation. In our previous study, we knocked out the *DCAF16* gene and performed untargeted global proteomics to identify proteins with increased abundance in *DCAF16* knockout cells as its potential endogenous substrates. Through this approach, we identified and validated SPIN4 as a substrate of DCAF16^10^. A potential limitation of this proteomics-based strategy is that it may not uncover additional E3 ligases that also regulate SPIN4 stability. For instance, although SPIN4 protein levels increase in *DCAF16* knockout cells, the contribution of other E3 ligases to SPIN4 degradation would remain uncharacterized unless they are directly targeted. As such, without prior knowledge of which E3 ligases to test, proteomics alone may be insufficient to reveal the complete regulatory network. In contrast, a CRISPR-based knockout screen provides an unbiased approach to systematically evaluate the contribution of most, if not all, E3 ligases to SPIN4 degradation.

To this end, we cloned SPIN4 into a dual-reporter vector encoding EGFP fused to SPIN4 and a separately translated mCherry (**Figure 3a**), and stably expressed SPIN4-EGFP in HEK293T cells. Fluorescence imaging confirmed that SPIN4-EGFP localized to the nucleus (**Figure S2**), consistent with its known subcellular localization and role as a histone methylation reader^10, 20^. Upon treatment with MG132 (a proteasome inhibitor) and MLN4924, we observed increased SPIN4-EGFP levels (**Figure 3b**), indicating that the overexpressed fusion protein is subject to degradation by the UPS, and more specifically, by the CRL pathway. These results suggest that SPIN4-EGFP likely retains its native conformation and serves as an ideal substrate system for the CRISPR screen to identify regulators of SPIN4 homeostasis.

**Figure 3.**
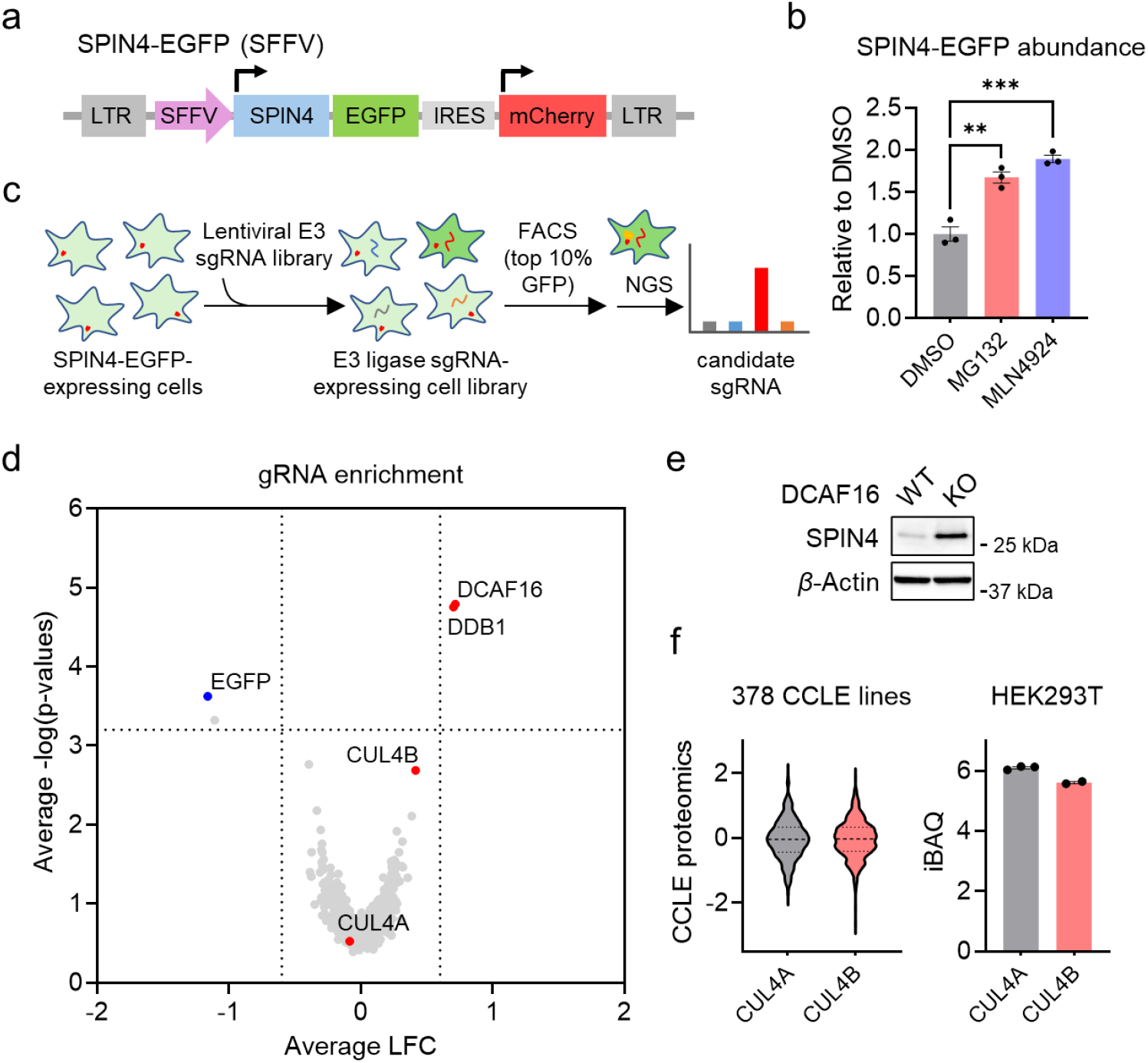
E3 ligase-focused CRISPR-Cas9 knockout screen to identify the E3 ligase required for SPIN4-EGFP degradation. **a**, The construct of SPIN4-EGFP. **b**, Quantification of SPIN4-EGFP fluorescence in HEK293T cells treated for 8 hours with 1 μM MG132 or 1 μM MLN4924. Data are presented as mean ± SEM (n = 3 biological independent samples). The statistical significance was assessed using unpaired two-tailed Student’s t-tests. **c**, Schematic overview of the CRISPR-Cas9 knockout screen workflow. **d**, Volcano plot showing results from the E3 ligase-focused CRISPR-Cas9 knockout screen for SPIN4-EGFP degradation (n = 3 biologically independent samples). *P* values were calculated by two-sided t test and adjusted using Benjamini-Hochberg correction for multiple comparisons. **e**, Western blot analysis of SPIN4 expression in MDA-MB-231 wild-type and *DCAF16* knockout cells. The result is representative of two experiments (n = 2 biologically independent samples). **f**, Published protein expression databases reveal CUL4A and CUL4B protein levels across Cancer Cell Line Encyclopedia (CCLE) cell lines and in HEK293T cells.

We next transduced Cas9 into SPIN4-EGFP-expressing HEK293T cells and applied the same focused sgRNA library for a CRISPR knockout screen. Following puromycin selection, we sorted the top 10% of cells with the highest GFP intensity and performed NGS to identify enriched sgRNA sequences, comparing them to those in the unsorted input population (**Figure 3c, Figure S3**, and **Table S2**). This screen identified DCAF16 and DDB1 as two top hits (**Figure 3d**). Since DCAF16 belongs to the CRL4 subfamily and interacts with the adaptor protein DDB1, our findings point to DCAF16-DDB1 as the sole E3 ligase complex responsible for SPIN4-EGFP degradation in HEK293T cells. Although no additional E3 ligases were uncovered, this result suggests a single regulatory mechanism governs SPIN4 degradation. To validate this, we examined endogenous SPIN4 expression in MDA-MB-231 *DCAF16* wild-type and knockout cells and observed elevated SPIN4 protein levels in the knockout cells (**Figure 3e**), consistent with previous reports^10, 15^.

Interestingly, data analysis revealed that sgRNAs targeting *CUL4B*, but not *CUL4A*, were enriched in the screen (**Figure 3d**). Although both CUL4A and CUL4B can function as scaffolds for DDB1 and substrate receptors, and proteomics databases indicate their expression levels are comparable^21, 22^ (**Figure 3f**), our results suggest that DCAF16 specifically depends on CUL4B for targeted degradation. We speculate that this selectivity arises from previously reported differences in subcellular localization: CUL4B contains a nuclear localization signal, whereas CUL4A is predominantly cytoplasmic^23^. Given that DCAF16 is a nuclear E3 ligase, our findings support the conclusion that it exclusively engages nuclear-localized CUL4B. This mechanistic insight is important for the precise modulation of DCAF16 function in future studies.

### CRISPR-Cas9 knockout screening reveals BAP1 as a DUB stabilizing SPIN4

Despite the physiological function of the DCAF16-SPIN4 axis remains unknown, SPIN4 acts as a reader of H3K4me3, a histone mark highly enriched at transcription start sites (TSSs) of actively transcribed genes^24^, suggesting this pathway may regulate gene transcription. Since protein homeostasis is often controlled not only by E3 ligases but also by DUBs, identifying potential DUBs for SPIN4 would help define the full regulatory pathway governing SPIN4 homeostasis. This would be critical for future studies aimed at understanding the physiological and pathological roles of SPIN4. To this end, we constructed another focused sgRNA library comprising 700 sgRNAs targeting 108 human

DUBs (six sgRNAs per DUB gene; **Table S1**) and performed a CRISPR pooled knockout screen to identify DUBs that stabilize SPIN4-EGFP (**Figure 4a** and **Figure S4**). Following puromycin selection, we sorted the bottom 10% of cells with the lowest GFP intensity, representing the population that lost DUB-mediated stabilization, and conducted NGS to identify enriched sgRNA sequences, comparing them to those in the unsorted input population (**Table S2**). This screen identified BRCA1-associated protein 1 (BAP1) as the top hit (**Figure 4b**).

**Figure 4.**
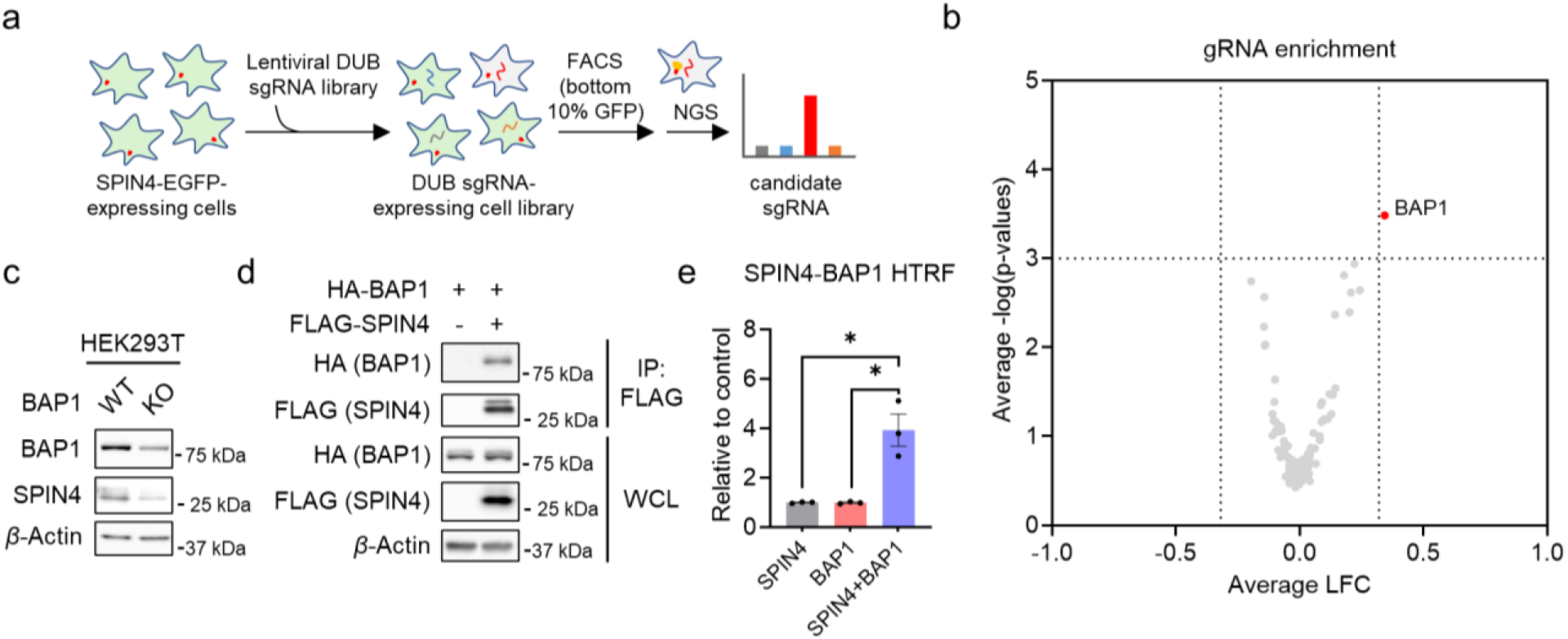
DUB-focused CRISPR-Cas9 knockout screen to identify the DUB required for SPIN4-EGFP stabilization. **a**, Schematic overview of the CRISPR-Cas9 knockout screen workflow. **b**, Volcano plot showing results from the DUB-focused CRISPR-Cas9 knockout screen for SPIN4-EGFP stabilization (n = 3 biologically independent samples). *P* values were calculated by two-sided t test and adjusted using Benjamini-Hochberg correction for multiple comparisons. **c**, Western blot analysis of SPIN4 and BAP1 expression in HEK293T wild-type and *BAP1* knockout cells. The result is representative of two experiments (n = 2 biologically independent samples). **d**, Co-immunoprecipitation analysis reveals an interaction between HA-BAP1 and FLAG-SPIN4. The result is representative of two experiments (n = 2 biologically independent samples). **e**, HTRF assay measuring the interaction between HA-BAP1 and FLAG-SPIN4 using anti-HA-d2 and anti-FLAG-Tb. Data are presented as mean ± SEM (n = 3 biological independent samples). The statistical significance was assessed using unpaired two-tailed Student’s t-tests.

We first validated this finding by generating *BAP1* knockout HEK293T cells and observed a decrease in SPIN4 expression (**Figure 4c**). Moreover, BAP1 and SPIN4 appear to physically interact, as demonstrated by both co-immunoprecipitation (**Figure 4d**) and homogeneous time-resolved fluorescence (HTRF) assays (**Figure 4e**). To further investigate this regulation, we performed quantitative global proteomics comparing wild-type and *BAP1* knockout HEK293T cells, and again observed reduced SPIN4 protein levels in the knockout cells (**Figure 5a** and **Table S3**). Additionally, an affinity purification-mass spectrometry (AP-MS) analysis of the SPIN4 interactome identified BAP1 as one of its interacting partners (**Figure 5b** and **Table S4**). The Spindlin family of proteins is largely conserved, and all members contain Tudor domains. We then asked whether SPIN1, the most well-studied Spindlin family member^25, 26^, can also be recognized and stabilized by BAP1. Both co-immunoprecipitation (**Figure S5a**) and HTRF (**Figure S5b**) assays indicate that BAP1 has minimal interaction with SPIN1. Moreover, global proteomics results reveal no difference in endogenous SPIN1 levels between wild-type and *BAP1* knockout HEK293T cells (**Figure S5c** and **Table S3**). Collectively, these data support that BAP1 interacts with and stabilizes SPIN4.

**Figure 5.**
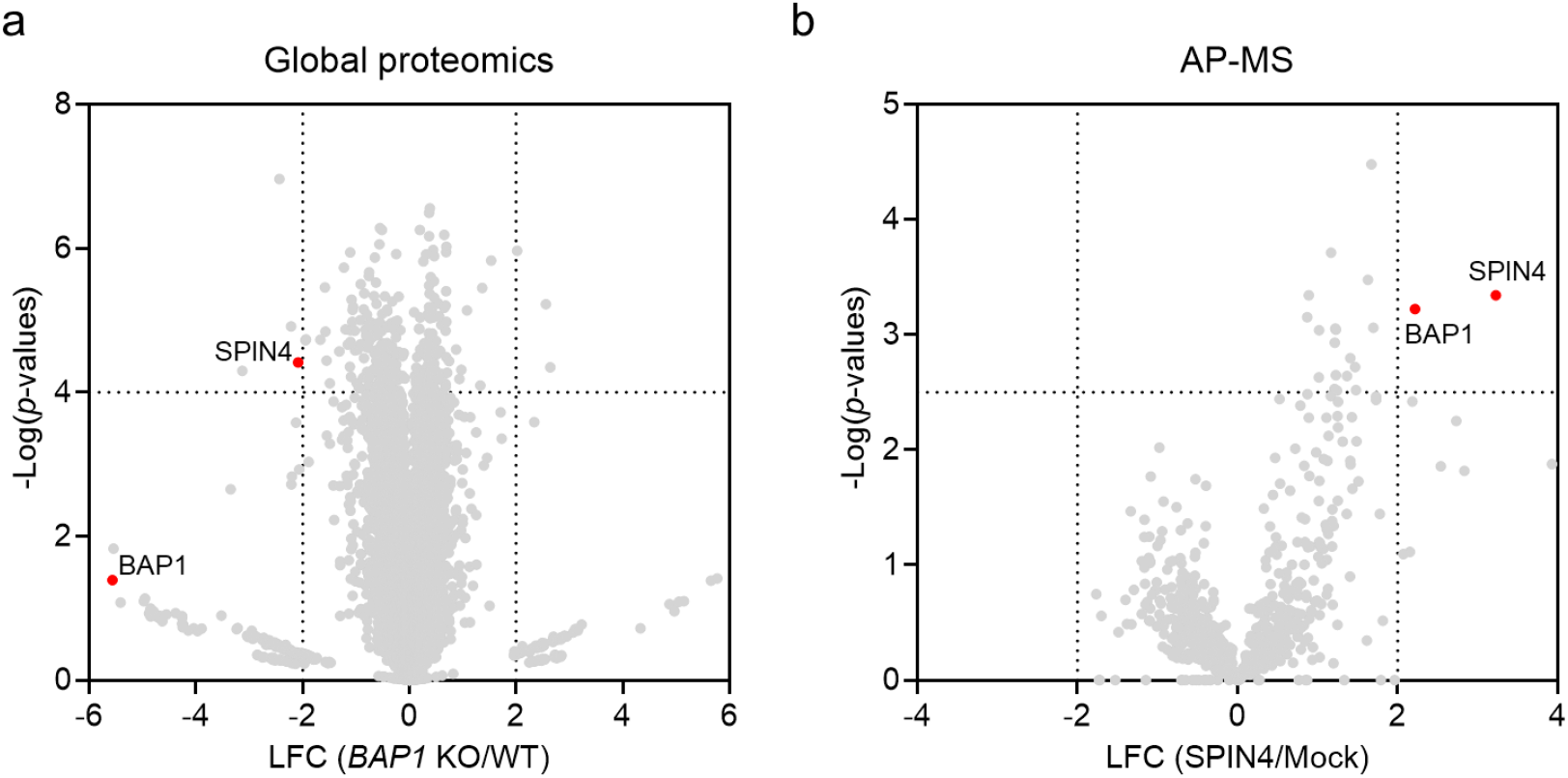
Proteomics analysis reveals that BAP1 interacts with and stabilizes SPIN4. **a**, Volcano plot showing global proteomic changes between wild-type and *BAP1* knockout HEK293T cells (n = 3 biological independent samples). *P* values were calculated by two-sided t test and adjusted using Benjamini-Hochberg correction for multiple comparisons. **b**, Volcano plot showing SPIN4 interactome analysis compared to mock-transfected cells (n = 3 biological independent samples). *P* values were calculated by two-sided t test and adjusted using Benjamini-Hochberg correction for multiple comparisons.

### BAP1 enzymatic activity is essential for SPIN4 stabilization

BAP1 is a cysteine protease that relies on its catalytic residue, Cys91, to mediate deubiquitination^27^. To determine whether BAP1’s enzymatic activity is essential for SPIN4 stabilization, we first treated HEK293T and 22Rv1 cells with a well-characterized BAP1 inhibitor, BAP1-IN-1 (**Figure 6a**), which blocks its catalytic function^28^. In both HEK293T and 22Rv1 cells, BAP1-IN-1 treatment led to a reduction in SPIN4 protein levels (**Figure 6b**). Using an HTRF assay, we further observed that BAP1-IN-1 treatment diminished the interaction between BAP1 and SPIN4 (**Figure 6c**). To directly test the importance of catalytic activity, we conducted quantitative proteomics in *BAP1*^C91S^ knock-in HEK293T cells. This mutant failed to restore SPIN4 expression, reinforcing the requirement of BAP1’s enzymatic activity for SPIN4 stabilization (**Figure 6d** and **Table S3**). Finally, analysis of proteomics data from 252 cancer cell lines in the Cancer Cell Line Encyclopedia (CCLE)^22^ revealed a positive correlation between BAP1 and SPIN4 protein levels (**Figure 6e**), supporting a potentially broader functional relationship between these two proteins across diverse cellular contexts. Given BAP1’s role as a tumor suppressor^27^, its regulation of SPIN4 may connect epigenetic control with tumorigenesis. Further studies are needed to elucidate the functional consequences of the SPIN4-BAP1 interaction in cancer and chromatin regulation.

**Figure 6.**
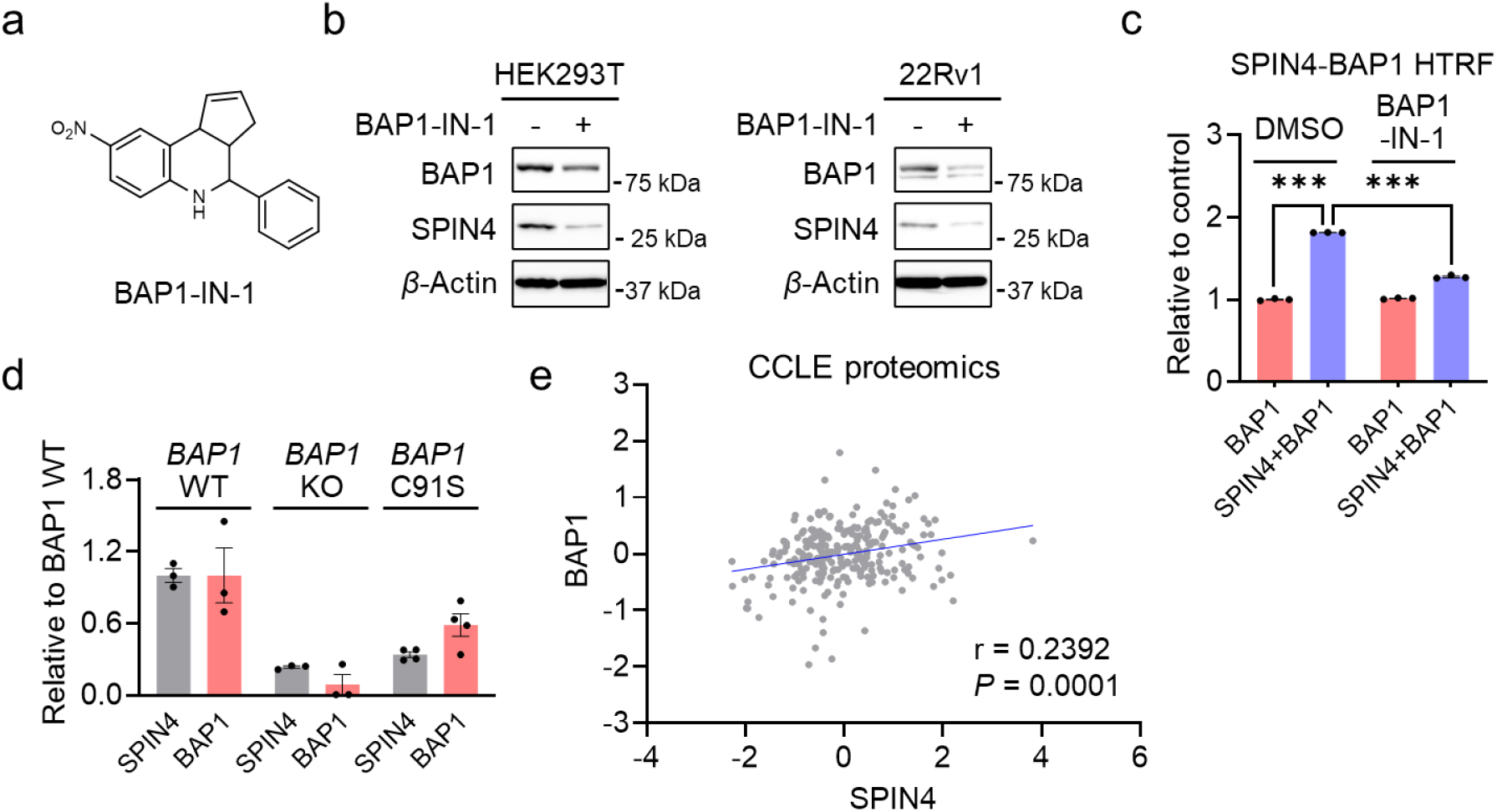
BAP1 enzymatic activity is essential for SPIN4 stabilization. **a**, Structure of BAP1-IN-1. **b**, Western blot analysis of SPIN4 and BAP1 expression in wild-type and *BAP1* knockout HEK293T and 22Rv1 cells treated with 10 µM of BAP1-IN-1 for 24 hours. The result is representative of two experiments (n = 2 biological independent samples). **c**, HTRF assay measuring the interaction between HA-BAP1 and FLAG-SPIN4 in the presence or absence of BAP1-IN-1 treatment (20 µM, 2 hours). Data are presented as mean ± SEM (n = 3 biological independent samples). The statistical significance was assessed using unpaired two-tailed Student’s t-tests. **d**, Global proteomics analysis comparing HEK293T wild-type, *BAP1* knockout, and *BAP1*^C91S^ knock-in cells revealed that BAP1-C91S failed to restore SPIN4 expression levels (n = 3 biological independent samples for wild-type and *BAP1* knockout cells, and n = 4 biological independent samples for *BAP1*^C91S^ knock-in cells). **e**, Correlation analysis of BAP1 and SPIN4 protein expression using data from the CCLE protein expression database.

## Conclusions

Through a CRISPR-Cas9 knockout screen, we identified SPIN4 as a substrate regulated by opposing pathways: stabilization mediated by BAP1 and degradation mediated by DCAF16. This regulatory balance governs SPIN4 protein abundance and may play a critical role in modulating its chromatin-associated functions. These findings suggest that precise control of SPIN4 stability may be integral to epigenetic regulation, with potential implications in disease contexts such as cancer.

## Supporting Information

Supplementary Figures 1-5, detailed materials and methods, and supplementary references.

Supplementary Table 1: List of sgRNA sequences for CRISPR knockout screens.

Supplementary Table 2: Average LFC and -log(p-values) of genes from CRISPR knockout screens.

Supplementary Table 3: Global proteomics comparing protein expression in HEK293T wild-type, *BAP1* knockout, and *BAP1*^C91S^ knock-in cells.

Supplementary Table 4: AP-MS analysis of HEK293T cells expressing FLAG-SPIN4.

## Acknowledgement

We gratefully acknowledge the support of the Falk Medical Research Trust Catalyst Award (X.Z.), NIH R35GM154945 (X.Z.), NIH T32GM149439 (A.S.), NIH R35GM146979 (L.W.), and International Institute for Nanotechnology Nano Scientist Program. We thank the Robert H. Lurie Comprehensive Cancer Center of Northwestern University for the use of the Flow Cytometry Core Facility.

## Notes

The authors declare no competing financial interest.

