## Supplementary Information for "CRISPR screen identifies BAP1 as a deubiquitinase regulating SPIN4 stability"

### Supplementary Figures

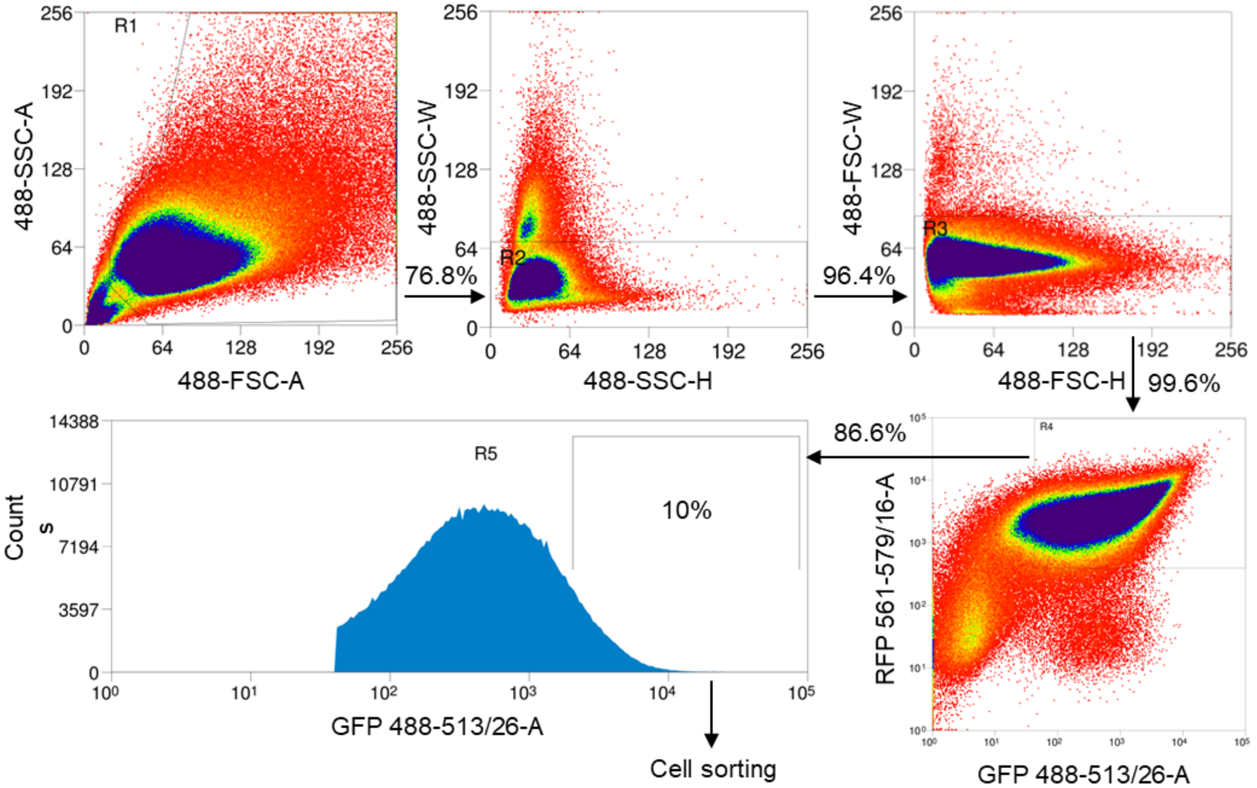


**Figure S1.** Gating strategy and fluorescence-activated cell sorting procedure for the CRISPR-Cas9 knockout screen to identify E3 ligases involved in dFKBP12-mediated FKBP12 degradation. Cells were first gated for singlets based on forward and side scatter. GFP-positive cells were then selected, and the top 10% of the GFP-expressing population was sorted and collected for downstream analysis.

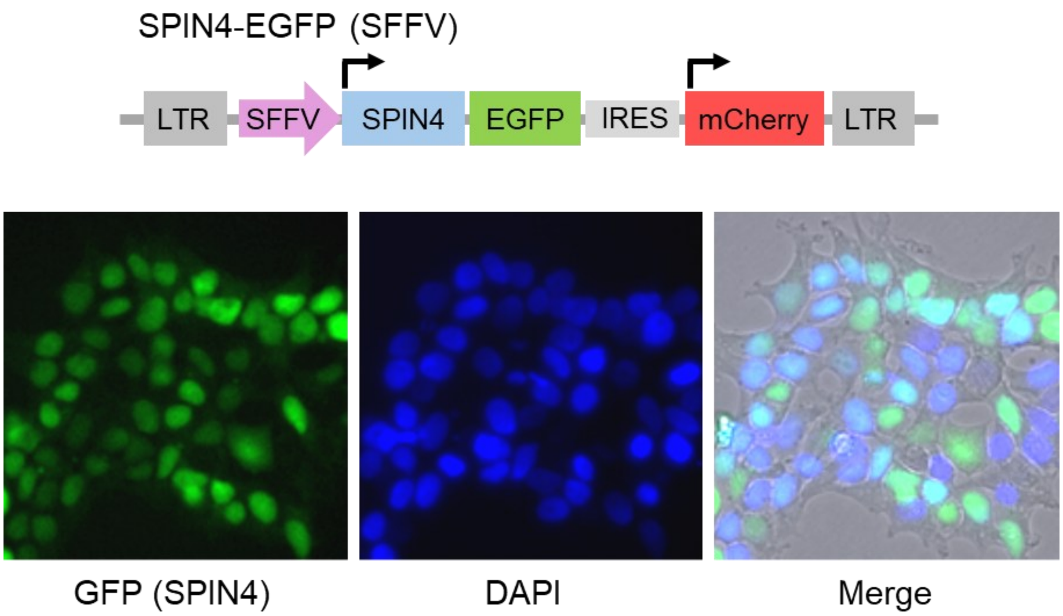


**Figure S2.** SPIN4-EGFP is primarily localized in the nucleus. DAPI (4′,6-diamidino-2-phenylindole) is a fluorescent dye used to stain cell nuclei.

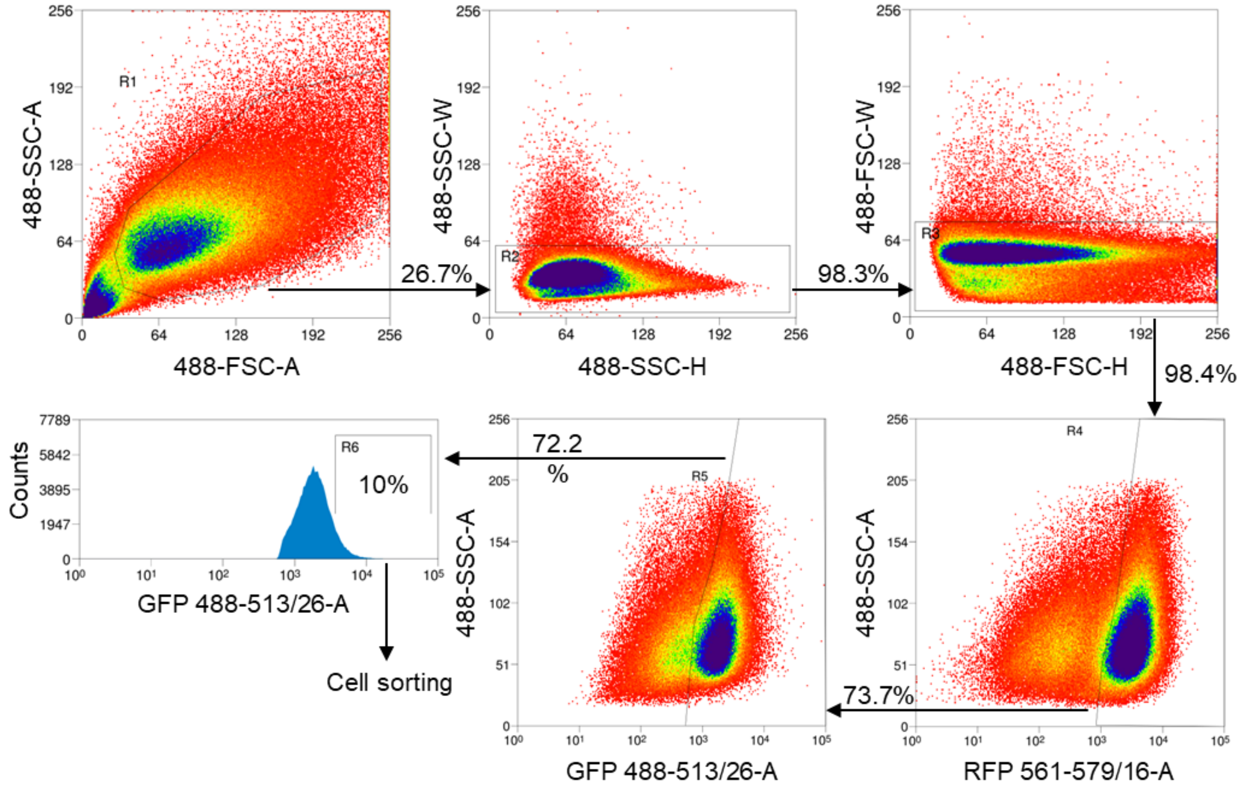


**Figure S3.** Gating strategy and fluorescence-activated cell sorting procedure for the CRISPR-Cas9 knockout screen to identify E3 ligases involved in SPIN4 degradation. Cells were first gated for singlets based on forward and side scatter. GFP-positive cells were then selected, and the top 10% of the GFP-expressing population was sorted and collected for downstream analysis.

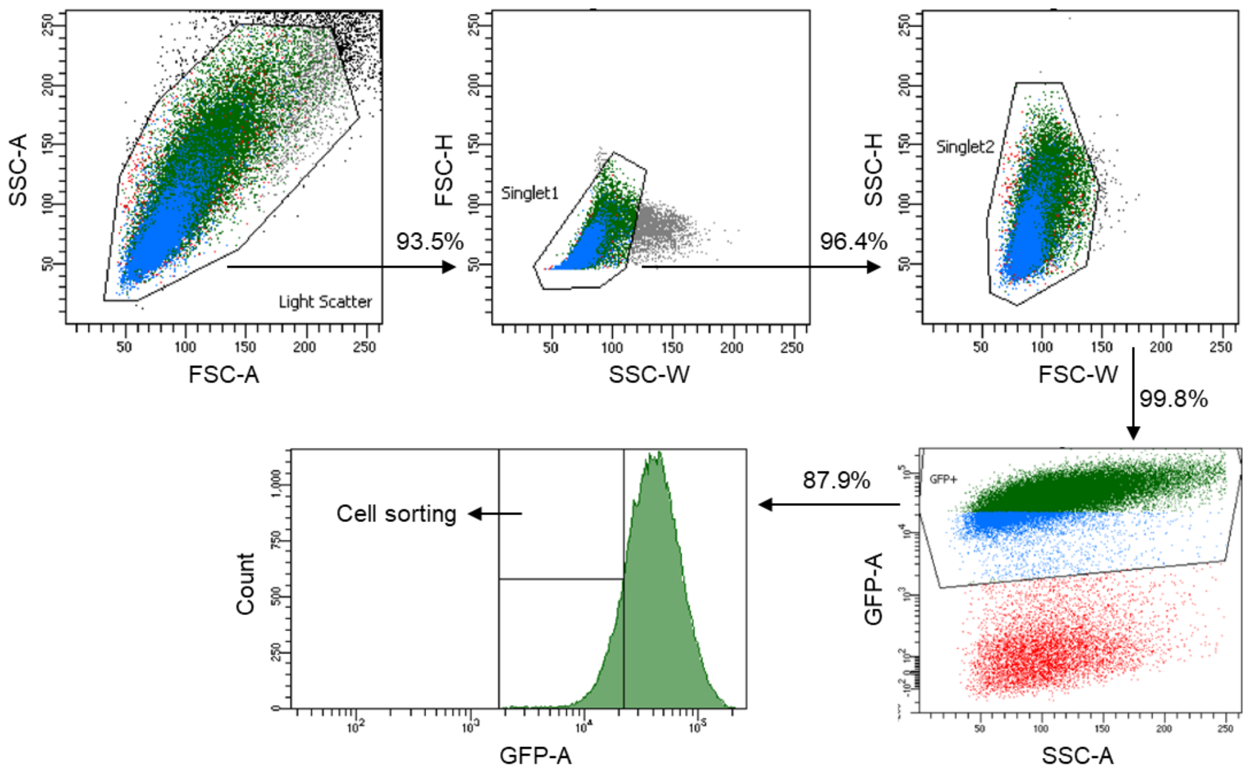


**Figure S4.** Gating strategy and fluorescence-activated cell sorting procedure for the CRISPR-Cas9 knockout screen to identify DUBs involved in SPIN4 stabilization. Cells were first gated for singlets based on forward and side scatter. GFP-positive cells were then selected, and the bottom 10% of the GFP-expressing population was sorted and collected for downstream analysis.

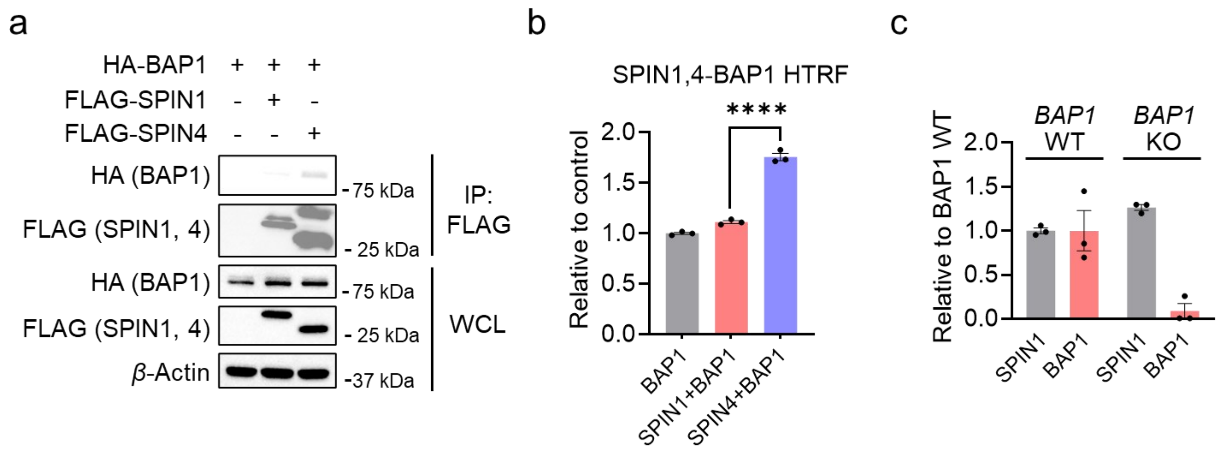


**Figure S5. BAP1 interacts predominantly with SPIN4 rather than SPIN1. a**, Co-immunoprecipitation analysis reveals an interaction between HA-BAP1 and FLAG-SPIN4, but not with SPIN1. The result is representative of two experiments (n = 2 biological independent samples). **b**, HTRF assay measuring the interaction between HA-BAP1 and FLAG-SPIN4 or FLAG-SPIN1 using anti-HA-d2 and anti-FLAG-Tb. Data are presented as mean ± SEM (n = 3 biological independent samples). The statistical significance was assessed using unpaired two-tailed Student's t-tests. **c**, Global proteomics analysis comparing HEK293T wild-type and *BAP1* knockout cells revealed that SPIN1 has similar expression levels (n = 3 biologically independent samples).

### Supplementary Materials and Methods

#### Reagents

The anti-FLAG HRP antibody (clone M2, cat. #A8592), anti-FLAG affinity gel (clone M2, cat. #A2220), anti-HA HRP antibody (clone 3F10, cat. #12013819001), and cOmplete protease inhibitor cocktail (cat. #11873580001) were purchased from Sigma-Aldrich. The anti-*β*-Actin HRP antibody (clone C4, cat. #sc-47778) was purchased from Santa Cruz Biotechnology. Cas9 endonuclease was purchased from Integrated DNA Technologies (cat. #1081061). FuGene 6 (cat. #E2692) transfection reagent and sequencing grade trypsin/Lys-C mix (cat. #V5071) were purchased from Promega. Puromycin (cat. #ant-pr-1) and blastcidin (cat. #ant-bl-05) were purchased from InvivoGen. MG132 (cat. #S2619) was purchased from Selleck Chemicals. MLN4924 (cat. #15217) and SLF (cat. #10007974) were purchased from Cayman Chemical. BAP1-IN-1 was purchased from MedChemExpress (cat. #HY-W327122). Enzyme-linked chemiluminescence (ECL) (cat. #32106), ECL plus (cat. #32132), BCA protein assay kit (cat. #23227), and Tandem Mass Tag (TMT) isobaric label reagent (cat. #90110) were purchased from Thermo Scientific. Polyethylenimine (PEI, MW 40,000, cat. #24765-1) was purchased from Polysciences, Inc.

#### Cell lines

HEK293T and MDA-MB-231 cells were obtained from ATCC and cultured in DMEM (Corning) supplemented with 10% FBS (v/v, Omega Scientific), L-Glutamine and penicillin-streptomycin antibiotic (Gibco). 22Rv1 cells were obtained from ATCC and cultured in RPMI 1640 (Corning) supplemented with 10% FBS (v/v, Omega Scientific), L-Glutamine and penicillin-streptomycin antibiotic (Gibco). All cell lines tested negative for mycoplasma contamination using the Universal Mycoplasma Detection Kit (ATCC, cat. #30-1012K).

#### Generation of CRSIPR-Cas9 *BAP1* knockout and *BAP1*$^{C91S}$ knock-in cells

To generate *BAP1* knockout cell lines in HEK293T and 22Rv1, three sgRNAs targeting *BAP1* were combined and used in electroporation for the delivery of Cas9-sgRNA RNP complex: BAP1 sgRNA #1 ACCCACCCTGAGTCGCATGA[1]; BAP1 sgRNA #2

AAGGTCTACCCCATTGACCA; BAP1 sgRNA #3 TTATGCCAAGTCCCCCATGC. BAP1-C91S knock-in cells were generated as previously described[2].

**Overexpression of BAP1, SPIN4 and SPIN1 in HEK293T cells**

HA tagged BAP1 plasmid (Addgene cat. #158109) was subcloned into pCDH-CMV-MCS-EF1-Blast vector using restriction sites NotI and NheI. FLAG tagged SPIN4 and SPIN1 plasmids were prepared as previously described[3]. Plasmids were used for PEI transient transfection of HEK293T cells, which were collected 48 hours post-transfection.

**Library preparation of sgRNAs**

sgRNAs targeting 680 human E3 ligases and 108 human DUBs were designed using CRISPick (Broad Institute), with six sgRNAs selected per gene. sgRNAs containing BsmBI restriction sites were synthesized as an oligonucleotide pool by Twist Bioscience. The sgRNA library was amplified by PCR and purified using the QIAquick gel extraction kit (Qiagen). The CRISPR knockout vector (LentiGuide-Puro, plasmid #52963) was obtained from Addgene. Pooled DNA was cloned into the CRISPR knockout vector using Esp3I (Thermo Scientific) and T7 ligase (New England Biolabs). The resulting construct library was transformed into ElectroMAX Stbl4 competent cells (Invitrogen) via electroporation using the Gene Pulser Xcell system (Bio-Rad). Transformed cells were cultured on Nunc Bioassay dishes (Thermo Scientific), and the construct library was extracted using the plasmid Maxi kit (Qiagen).

**CRISPR-Cas9 knockout screen**

Lentivirus containing the sgRNA library targeting human E3 ligases or DUBs was transduced into FKBP12-EGFP or SPIN4-EGFP and Cas9 (pLX_311-Cas9, Addgene plasmid #96924)-expressing HEK293T cells at a multiplicity of infection (MOI) of 0.3. 24 hours post-transduction, cells were treated with 2 μg/mL puromycin for 4 days. Following puromycin selection, cells were either treated with 1 μM dFKBP12 for 8 hours (FKBP12 screen) or directly proceeded to the next step (SPIN4 screen). Cells were then sorted by BD FACSAria III based on GFP fluorescence, selecting the top 10% for the E3 screen and bottom 10% for the DUB screen. Total DNA was extracted using the NucleoSpin Blood Mini Kit (MACHEREY-NAGEL). sgRNAs were amplified by PCR using a mixed P5 primer and a P7 primer containing an i7 index sequence. sgRNA abundance was quantified via Illumina MiSeq sequencing. Read counts for sgRNAs targeting each gene were used to calculate fold changes and *p*-values.

**Homogenous time resolved fluorescence assay**

HEK293T cells overexpressing HA-BAP1 or co-expressing HA-BAP1 with FLAG-SPIN4 or FLAG-SPIN1 were collected and lysed by sonication in PBS containing 0.1% BSA. Protein concentration was quantified using BCA assay (cat. #23227, Thermo Fisher Scientific) and normalized to 2 mg/mL. In a white, low-volume 96-well HTRF plate (cat. #NC1286225, Cisbio), 10 µL of cell lysate was added per well, followed by 5 µL of diluted anti-HA-d2 (cat. #610HADAF, Cisbio) and 5 µL of anti-FLAG-Tb antibodies (cat. #61FG2TLF, Cisbio), then incubated at room temperature for 1 hour. Fluorescence signals were measured using a CLARIOstar Plus microplate reader (BMG LABTECH) at 620 nm and 665 nm. The average background fluorescence from lysates without antibodies was subtracted from the sample wells, and the fluorescence ratio of 665 nm to 620 nm was calculated.

**Western blot analysis**

Cells were lysed in NP-40 lysis buffer (25 mM Tris-HCl, pH 7.4; 150 mM NaCl; 10% glycerol; 1% Nonidet P-40) supplemented with cOmplete protease inhibitor cocktail and sonicated. Protein concentrations were quantified using the BCA assay. Samples were then mixed with 4× Laemmli sample buffer and incubated at 95 °C for 5 minutes. Protein samples were loaded onto 4-20% Novex Tris-Glycine mini gels (Thermo Scientific) and transferred onto 0.2 µm polyvinylidene fluoride (PVDF) membranes (Bio-Rad). Membranes were blocked with 5% nonfat milk in TBST buffer for 1 hour at room temperature. Anti-FLAG, anti-HA, and anti-β-actin HRP-conjugated antibodies were diluted 1:5000 in blocking solution, while anti-BAP1 and anti-SPIN4 antibodies were diluted 1:1000. All antibodies were incubated overnight at 4°C. Prior to secondary antibody incubation (1:5000 dilution for 1 hour at room temperature), membranes were washed three times for 5 minutes each with TBST. Following secondary antibody incubation, membranes were washed three times for 5 minutes each with TBST, then incubated with ECL western blotting detection reagent to develop the chemiluminescence signal. Blots were imaged using the ChemiDoc MP system (Bio-Rad).

### Immunoprecipitations

HEK293T cells overexpressing HA-BAP1 or co-expressing HA-BAP1 with FLAG-SPIN4 or FLAG-SPIN1 were lysed in NP-40 lysis buffer (25 mM Tris-HCl, pH 7.4; 150 mM NaCl; 10% glycerol; 1% Nonidet P-40) supplemented with cOmplete protease inhibitor cocktail and sonicated. Lysates were centrifuged to collect the supernatant, and protein concentrations were quantified and normalized to 2 mg/mL. Normalized lysate was added to 30 µL of FLAG affinity gel beads and rotated at 4°C for 2 hours. The beads were then washed eight times with immunoprecipitation washing buffer (0.2% NP-40, 25 mM Tris-HCl, pH 7.4; 150 mM NaCl). After the final wash, the beads were incubated with 2× Laemmli sample buffer and heated at 95°C for 5 minutes. The supernatant was collected and used for western blot analysis.

### Global proteomics analysis

Cells were lysed in PBS supplemented with cOmplete protease inhibitor cocktail and sonicated, followed by protein quantification and normalization to 2 mg/mL using the BCA assay. From this, 50 µL of protein lysate was mixed with 50 µL of 12 M urea and 5 µL of 200 mM DTT, then incubated at 65°C for 15 minutes. Samples were alkylated by adding 5 µL of 400 mM iodoacetamide and incubated in the dark at 37°C for 30 minutes. PBS was added to dilute the urea concentration to 2 M, followed by the addition of trypsin for digestion at 37°C for 18 hours. For TMT labeling, 35 µL of each digested sample was mixed with 9 µL of acetonitrile and TMT tags, and incubated at room temperature for 1 hour. The reaction was quenched by adding 6 µL of 5% hydroxylamine and incubating at room temperature for 15 minutes, then acidified with 2.5 µL of formic acid. All samples were pooled and desalted using Sep-Pak C18 cartridge (Waters). Desalted samples were dried and subjected to fractionation using the High-pH Reversed-Phase Peptide Fractionation Kit (Thermo Fisher), resulting in 10 distinct fractions.

Peptide fractions were analyzed by liquid chromatography-tandem mass spectrometry on an Orbitrap Eclipse Tribrid Mass Spectrometer coupled to a Vanquish Neo UHPLC system. Peptides were loaded onto an EASY-Spray C18 HPLC column (2 µm particle size, 75 µm inner diameter, 150 mm length) and eluted at 0.25 µL/min using the following gradient: 5% buffer B (80% acetonitrile, 0.1% formic acid) in buffer A (water, 0.1% formic acid) from 0-15 minutes, increased to 35% buffer B from 15-155 minutes, then ramped to 100% buffer B from 155-180 minutes. The nano-LC electrospray ionization source was set to 1.5 kV. In MS1, data were collected at a resolution of 120,000 with an *m/z* range of 375-1600, RF lens set to 30%, standard AGC target, and auto maximum injection time. In MS2, precursor ions were isolated by quadrupole (0.7 *m/z* isolation window) and fragmented via HCD in the ion trap (collision energy 32%; standard AGC target; maximum injection time 35 ms). Following each MS2 scan, synchronous precursor selection (SPS) enabled selection of up to 10 MS2 fragment ions for MS3 analysis, where precursors were fragmented by HCD and analyzed in the Orbitrap (collision energy 55%; AGC target 250%; maximum injection time 200 ms; resolution 50,000). RAW data were acquired in Xcalibur (v4.5.445.18) and analyzed using Proteome Discoverer 2.5.

**Affinity purification coupled with mass spectrometry (AP-MS)**

Samples were prepared and subjected to FLAG enrichment as described above for immunoprecipitation. After the final wash of the affinity beads with immunoprecipitation buffer, the beads were washed once more with PBS and pelleted by centrifugation. The PBS was removed, and the beads were resuspended in 8 M urea in PBS and incubated at 65 °C for 10 minutes. The beads and supernatant were then transferred to a BioSpin column (Bio-Rad) to elute proteins. The eluate was reduced with 5 µL of 200 mM DTT and incubated at 65 °C for 15 minutes, followed by alkylation with 5 µL of 400 mM iodoacetamide at 37 °C for 30 minutes in the dark. Urea in the mixture was diluted to 2 M by adding PBS, and proteins were digested with trypsin for 18 hours at 37°C. To each sample, 5 µL of TMT tags were added and incubated at room temperature for 1 hour. The reaction was quenched by adding 6 µL of 5% hydroxylamine, incubated at room temperature for 15 minutes, and acidified with 4 µL of formic acid. All samples were pooled, desalted using a Sep-Pak C18 cartridge, and dried in a SpeedVac concentrator before mass spectrometry analysis as described above.

**Statistical analysis**

Quantitative data are presented as scatter plots with the mean and standard error of the mean (SEM) shown as error bars. Comparisons between two groups were performed using an unpaired two-tailed Student's t-test. Statistical significance is indicated as follows: *$P < 0.05$, **$P < 0.01$, ***$P < 0.001$, and ****$P < 0.0001$.
